# Single-Cell RNA-seq Reveals a Subpopulation of Cells Underlying β Cell Expansion in the Postnatal Islets

**DOI:** 10.1101/303263

**Authors:** Jingli A. Zhang, Chunyan Gu, Derek K. Smith, Monica K. Beltran, Noelyn Kljavin, Hai Ngu, Rowena Suriben, Jeremy Stinson, Zora Modrusan, Andrew S. Peterson

**Author notes:** Present address: Amgen Inc, South San Francisco, CA 94010, USA. Present address: Corvus Pharmaceuticals, Burlingame, CA 94010, USA. Present address: NGM Biopharmaceuticals, South San Francisco, CA 94080, USA.

## Abstract

Pancreatic β cells undergo significant expansion and maturation during human and rodent postnatal development. Here, we used single-cell RNA-seq to characterize gene expression patterns at various stages of mouse islet cell development and uncovered a population of cells that is most abundant during the early postnatal period. This cell population lacks expression of FLTP and expresses PDGF receptors. Each of these conditions have previously been associated with proliferative capacity in β cells suggesting that we have identified the proliferative competent of β cell mass expansion. The subpopulation co-express many endocrine lineage-specific genes and exhibits a downregulation of genes associated with mitochondrial oxidative phosphorylation and global protein synthesis. It has upregulated activity of genes in the Wnt, Hippo, PDGF, and Notch pathways and has a significantly higher proliferation potential than the more mature β population. We show that activity of the Notch pathway is required in postnatal β cell expansion where it serves to maintain an undifferentiated endocrine state in the polyhormonal cell population. Collectively, our study identifies a proliferative, progenitor-like cell subpopulation in the postnatal islet as the source of postnatal β cell expansion.

## Introduction

Pancreatic β cells naturally adapt, through increased insulin production and compensatory β cell proliferation, to meet high insulin demand in physiological and pathological circumstances, including growth, pregnancy, obesity and injury (Dor et al. 2004; Brennand et al. 2007; Nir et al. 2007; Teta et al. 2007; Meier et al. 2008). Diabetes mellitus results when adaptation fails to produce sufficient functional β cell mass. While type 1 diabetes (T1D) is caused by autoimmune destruction of β cells, progression to type 2 diabetes (T2D) includes an initial compensatory expansion of β cell mass via increased β cell replication, in response to peripheral insulin resistance, followed by a progressive loss of functional β cells and overt hyperglycemia (Heit et al. 2006; Rahier et al. 2008; Talchai et al. 2009). Obesity is a major cause of peripheral insulin resistance; however, only a third of obese adults develop T2D, implying that defects in adaptive mechanisms are likely to feature in the pathogenesis of T2D (Linnemann et al. 2014). Harnessing the adaptive responses of β cells has been proposed as an attractive approach to treat both T1D and T2D (Uchida et al. 2005; Butler et al. 2007; Nir et al. 2007; Linnemann et al. 2014; Cheng et al. 2017).

After birth, replication of pre-existing, insulin expressing β cells is the predominant means for β cell mass increase in both humans and rodents (Dor et al. 2004; Brennand et al. 2007; Nir et al. 2007; Teta et al. 2007; Gregg et al. 2012). Based on this, expansion of adult human β cells *in vitro* has been explored as a replacement cell therapy for diabetes. However, full expression of β cell features and functional activities in *in vitro* culture is a major hurdle still to be overcome (D’Amour et al. 2006; Russ et al. 2009; Bar et al. 2012; Lenz et al. 2014). Therefore, an understanding of the extrinsic and intrinsic regulations governing β cell regeneration, as well as demarcation of the molecular differences between proliferating and functional β cells could help identify new therapeutic targets for diabetes prevention and treatment.

During physiological growth, β mass expansion is dependent upon PDGF signaling and has a characteristic time course with proliferation peaking in the early postnatal period and gradually declining thereafter to adulthood (Teta et al. 2005; Krishnamurthy et al. 2006; Meier et al. 2008; Chen et al. 2011). Although β cells have long been considered to form a homogeneous population that broadly retains regeneration potential (Brennand et al. 2007; Teta et al. 2007), recent studies have indicated heterogeneity in the proliferative capacity of this population, both in humans and rodents (Bader et al. 2016; Dorrell et al. 2016; Wang et al. 2016). In particular, FLTP, a Wnt/planar cell polarity (PCP) effector, acts as a biomarker that distinguishes proliferation-competent from mature β cells. *Fltp^−^Nkx6.1^+^* β cells, which are more evident in postnatal animals and during pregnancy, have poorer glucose sensing and insulin secretion, and show marked higher proliferative capacity, compared to *Fltp^+^Nkx6.1^+^* β cells (Bader et al. 2016).

To explore β cell heterogeneity during the postnatal period and gain insight into differences between proliferating and mature β cells, we profiled the gene expression patterns of islet cells at multiple developmental stages (E18.5, P1.5, P4.5, P7.5 and adult) using single-cell RNA-seq (Gutierrez et al. 2017). This allowed us to identify a *Fltp^−^, Nkx6.1^+^* population that is most apparent at P7.5. Differential gene expression analyses distinguish this cell population from *Nkx6.1^+^* β cells despite the fact that both populations express transcription factors, such as *Isl1, Nkx2.2, Nkx6.1, Pax6, Pdx1* and *Mafb*, that are characteristic of β cells. Importantly, we show that during the postnatal period of rapid β cell mass expansion this cell population displays significantly higher proliferation potential than more mature β cells and expresses both receptors and ligands of the PDGF pathway, and indicating that an autocrine growth mechanism may be involved in postnatal β cell expansion. Relative to β cells and α cells of all ages studied, the *Fltp, Nkx6.1^+^\* population expresses low levels of both insulin and glucagon transcripts and very low levels of genes associated with protein synthesis and mitochondrial oxidative phosphorylation, a pattern seen in a number of stem or progenitor cell populations (Signer et al. 2014; Llorens-Bobadilla et al. 2015; Wanet et al. 2015; Blanco et al. 2016). This cell population also expresses receptors and ligands of the Wnt, Hippo and Notch signaling pathways as well as their corresponding downstream targets suggesting that a broad autocrine program is involved in β cell expansion. We further demonstrate that Notch signaling maintains these cells in an undifferentiated state and is required for β cell expansion. Taken together, our studies identify a subpopulation of cells in the islets at P7.5 that are maintained in an undifferentiated and proliferative progenitor-like state through autocrine or paracrine regulation and that drive postnatal β cell expansion.

## Results

### β cell proliferation peaks one week after birth

While total β cell mass increases continuously from birth until adulthood, there is significantly more proliferation at age P7.5 relative to P1.5, P21, and young adult (3 months old) (Fig. 1A, B). To understand the gene expression changes that might promote this increase in β cell proliferation during the early postnatal period, we used fluorescence activated cell sorting (FACS) followed by RNA sequencing (RNA-seq) to define the transcriptomes of eGFP-positive β cells from E18.5, P1.5, P7.5, P21, and young adult (3-month old) *MIP1-eGFP* mice (Hara et al. 2003). Pairwise comparisons between consecutive ages, as well as between E18.5 and young adult, and P7.5 and young adult, identified 2,310 differentially expressed transcripts (Supplemental Table S1). Unsupervised hierarchical clustering partitioned these genes into four distinct clusters (Fig. 1C). Corresponding well with the gradual maturation of glucose-stimulated insulin secretion of β cells after birth (Blum et al., 2012), genes involved in hormone processing and secretion were upregulated with increasing age, including *G6pc2, Ins1/2, Pam, Pcsk1, Pparg, Scg2, Scg5, Scl2a2*, and *Ucn3* (Cluster I); in parallel, genes associated with tissue and organ development such as *Egr1, Fos, Fosb, Jun, Junb, Sox9*, and several Krüppel-like factors were gradually downregulated (Cluster III) (Fig. 1C; Supplemental Table S1).

**Figure 1.**
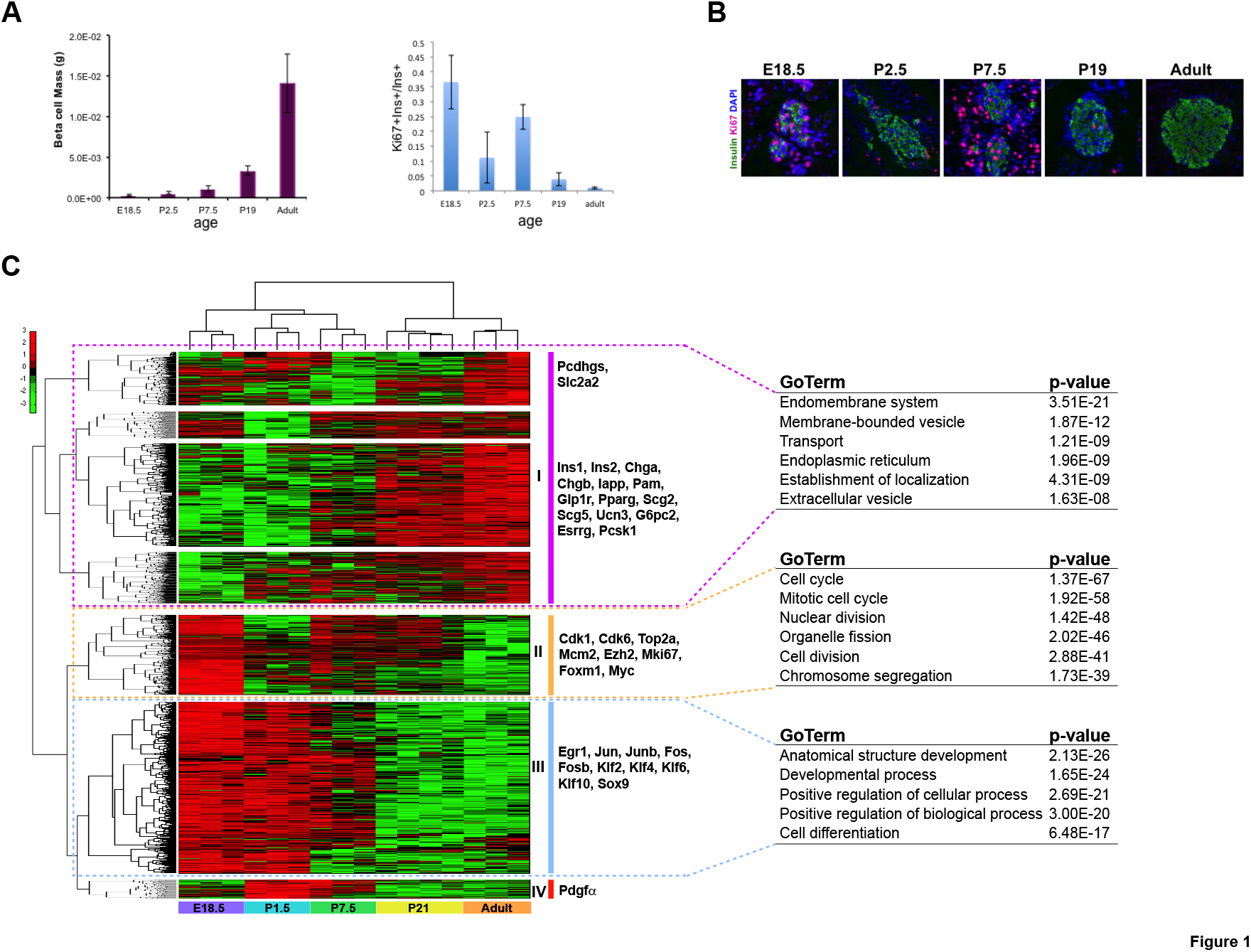
Increase in β cell proliferation during postnatal period. (A) Quantification of β cell mass of pancreas (left); quantification of β cell KI67 labeling (right) at five ages. (B) Immunofluorescence staining of INS1 and KI67 in islets at five ages. (C) Heatmap showing differentially expressed genes (FDR < 0.01) in FACS sorted *MIP1-GFP^+^* β cells across five ages: E18.5, P1.5, P7.5, P21 and adult (columns). There were three biological replicates each stage, except for P21 (four replicates). Genes are clustered into 4 groups based upon the patterns, and some representative genes are listed next to each block. Data (log_2_(RPKM+0.0001)) were standardized along the rows such that the mean is 0 and the standard deviation is 1. Green and red colors correspond to minimum and maximum expression, respectively. Selected, top significantly enriched GO terms for genes in the first three clusters (cluster I, cluster II and cluster III) are listed next to the heatmap.

On the other hand, mirroring the proliferation pattern of β cells during the initial period after birth, there was a biphasic pattern of cell division-associated genes with significantly elevated expression at E18.5, P7.5, and P21, such as *Cdk1, Cdk6, Ezh2, Foxm1, Mcm2, Mki67, Myc*, and *Top2a* (Cluster II) (Fig. 1C; Supplemental Fig. S1; Supplemental Table S1). Interestingly, we also found a small group of genes with peak expression immediately following birth at P1.5 and preceding the peak of cell division at P7.5 (Cluster IV) (Fig. 1C; Supplemental Fig. S1; Supplemental Table S1). Amongst these genes we could detect *Pdgfa*, a gene of particular interest since PDGF signaling, which acts through *Ezh2* induction, is required for the proliferative capacity of pancreatic β cells in young mice (Chen et al. 2011). The upregulation of *Pdgfa* in anticipation of the proliferative surge at P7.5 and the corresponding induction of *Ezh2* that parallels proliferation is consistent with the prior conclusion that this pathway drives postnatal β cell proliferation.

### Single-cell RNA-seq identifies four clusters of cells in adult islets

We next explored the use of single-cell RNA-seq (scRNA-seq) as a method for defining the transcriptional profiles of individual cells in the postnatal islet (Xin et al. 2016). As a first step we used young adult islet cells from C57BL/6 mice to evaluate the potential utility of this approach. Purified islets were handpicked from 3-month old C57BL/6 mice, dissociated into single cells, and captured using the Fluidigm C1 integrated fluidic circuit for RNA-seq (Chen et al. 2017) (Materials and Methods). In total, 159 single cell transcriptome datasets from two independent experiments passed the selection filters designed to exclude dying cells and doublets (Materials and Methods). To measure the molecular similarity across individual cells, we calculated the cell-to-cell pairwise Pearson correlation coefficients. The correlation analysis excluded genes in the bottom 10% variance bin, as well as any genes with less than 0.1 RPKM in all selected cells. We also excluded the 15 highest expressed cell-type specific genes, including those encoding the cell-type specific endocrine hormones, from the coefficient calculation, reasoning that these highly expressed genes could overwhelm and obscure the recognition of important cell subtypes or sub-states (Materials and Methods). Although the exclusion of the top 15 cell-type specific genes led to a decrease in pairwise correlation coefficients in general, it did help us uncover hidden subpopulations that shared some hormone signatures with other subpopulations. The hierarchical clustering of the resulting correlation coefficients (Fig. 2A, top) revealed three distinct endocrine cell types that, when compared with the expression of endocrine hormone genes (*Gcg, Iapp, Ins1, Ins2, Pyy, Ppy*, and *Sst)*, appeared consistent with the canonical view of the major endocrine islet cell types: β cells (*Gcg*^low^, *Ins1*^high^, *Ins2*^high^, and *Sst*^low^), a cells (*Gcg*^high^, *Ins1^1^™*, *Ins2*^low^, and *Sst*^low^), and 5 cells (*Gcg*^low^, *Ins1*^low^, *Ins2*^low^, and *Sst*^high^) (Fig. 2A, bottom). Additionally, five cells were identified as PP cells with *Ppy* representing the predominant hormone gene; however, they did not independently cluster and were instead mixed within α and δ populations (Supplemental Table S2). Interestingly, most α cells and δ cells exhibited significant expression of *Ppy* and/or *Pyy*, while β and δ cells expressed *Iapp* (Fig. 2A, bottom).

**Figure 2.**
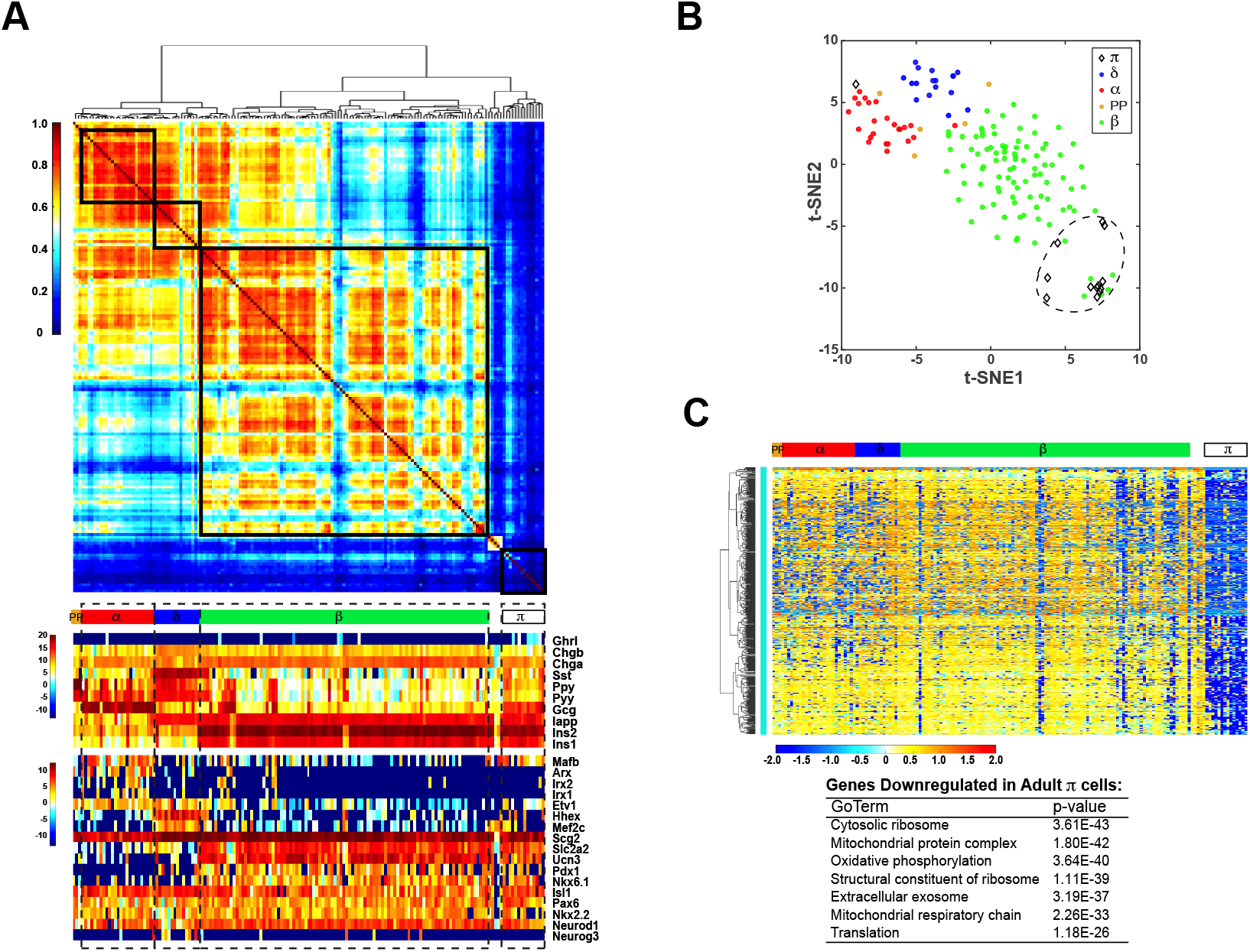
Single-cell transcriptome signature of adult mouse islets (A) Up: Correlation coefficient cluster plot partitions adult islet cells into δ, α, β and π (polyhormonal) subpopulations (five PP cells were identified based on the dominant expression of *Ppy* compared to other hormones, but not enough to form a separated cluster; five cells between β group and π group were treated as unknown and excluded as Itgax was detected in these cells in addition to hormones). Color scale corresponds the absolute correlation coefficient of each pair of cells; Bottom: Heatmaps showing expression patterns of selected cell type specific hormones and transcription factors. Color scale corresponds the expression level (log_2_(RPKM+0.0001)). (B) t-SNE plot of adult islet scRNA-seq data, based on 1327 differentially expressed genes (FDR < 0.01). Each dot or diamond represents one cell, and cell identity is indicated by color or shape. (C) Heatmap showing a group of genes uniquely downregulated in π population (Figure S2A). Data (log_2_(RPKM+0.0001)) are standardized along each gene (row), and color scale corresponds the relative expression level. Selected, top significant enriched GO terms are listed in the table beneath the heatmap.

Sporadic bihormonal cells of indeterminate identity were found within each cluster of canonical, clearly-identifiable cell types. We were unable to determine whether these cells represent a unique non-canonical cell type, a transitional cell state or the incomplete removal of cell doublets. In contrast to these sporadic cells, we identified a fourth cluster of cells that showed dramatically reduced hormone transcript expression (*Gcg*^low^, *Ins1/2*^low^, and *Sst*^low^) (14 of 159 cells, Fig. 2A). Due to their unique gene expression pattern it was apparent that these were not the result of capturing cell aggregates. These cells showed striking similarity to the PDGF expressing cells described below, so to be consistent with our designation of the proliferative cell compartment described below, we referred to these cells as π cells (Fig. 2; Supplemental Fig. S2).

### Differentially expressed genes in adult islets

To confirm known cell-type specific gene expression and to identify novel molecular markers for islet cell types, we performed differential expression analysis on selected young adult islet cells using Monocle. Using this approach we identified 1,327 differentially expressed genes (FDR < 0.01) (Materials and Methods). Unsupervised hierarchical clustering of these genes identified enrichment of the well-established markers *Nkx6.1, Pdx1, Scl2a2*, and *Unc3* in the β cell the cluster and enrichment of *Arx, Mafb, Irx1* and *Irx2* in the α cell cluster (Petri et al. 2006; Romer and Sussel 2015). We further confirmed enrichment of *Hhex*, a recently discovered δ cell-specific transcription factor, and *Pdx1* in the δ cell population (Zhang et al. 2014) (Fig. 2A; Supplemental S2A; Supplemental Table S2).

In addition to these established genes, we identified a couple of novel cell-type specific markers, such as *Mef2c* for δ cells and *Etv1* for both α cells and δ cells (Fig. 2A; Supplemental Table S2). Surprisingly, the endocrine progenitor factor *Neurog3* was detected more often in the δ cell population (~50% of δ cells) relative to β cell, α cell, and π cell populations (< 5% β cells, α cells, and π cells). To further investigate the relationship between the four cell populations in adult mouse islets, we performed t-Distributed Stochastic Neighbor Embedding (t-SNE) analysis (Materials and Methods). As expected, t-SNE clustered the majority of cells into four distinct groups (β, α, δ, and π) with the π group closer to the β group than either the α or δ groups (Fig. 2B). Interestingly, several β cells clustered within the π group implying that these cells were likely either intermediate cells or π cells that were misidentified as β cells due to the molecular similarity of two cell types.

### P7.5 islets are enriched with two types of polyhormonal cells

To identify the cell types associated with proliferation in the early postnatal islet and unravel the regulatory mechanisms underlying this process, we handpicked islets from P7.5 *MIPl-eGFP* mice and generated 201 scRNA-seq datasets from three independent experiments (Materials and Methods). Applying the same approach used to analyze adult scRNA-seq data we provisionally identified six cell types in P7.5 islets based on the correlation matrix dendrogram and expression pattern of hormone signatures (Fig. 3A). As expected, approximately half of the cells could be categorized as β cells (*Gcg*^low^, *Ins1*^high^, *Ins2*^high^, and *Ss*^low^), while one-fourth resembled *α* cells (*Gcg*^high^, *Ins1*^low^, *Ins2*^low^, and *Sst*^low^). Interestingly, the relative expression of *Pyy* in P7.5 β cells is significantly higher than in young adult β cells (Fig. 3A). Although some cells expressed *Sst*, most expressed significant levels of *Gcg, Ins1, Ins2* or other hormones and did not have a sufficiently unique pattern of gene expression to cluster as a distinct δ group, which possibly reflects the lineage flexibility of postnatal *Sst*-positive cells (Chera et al. 2014).

**Figure 3.**
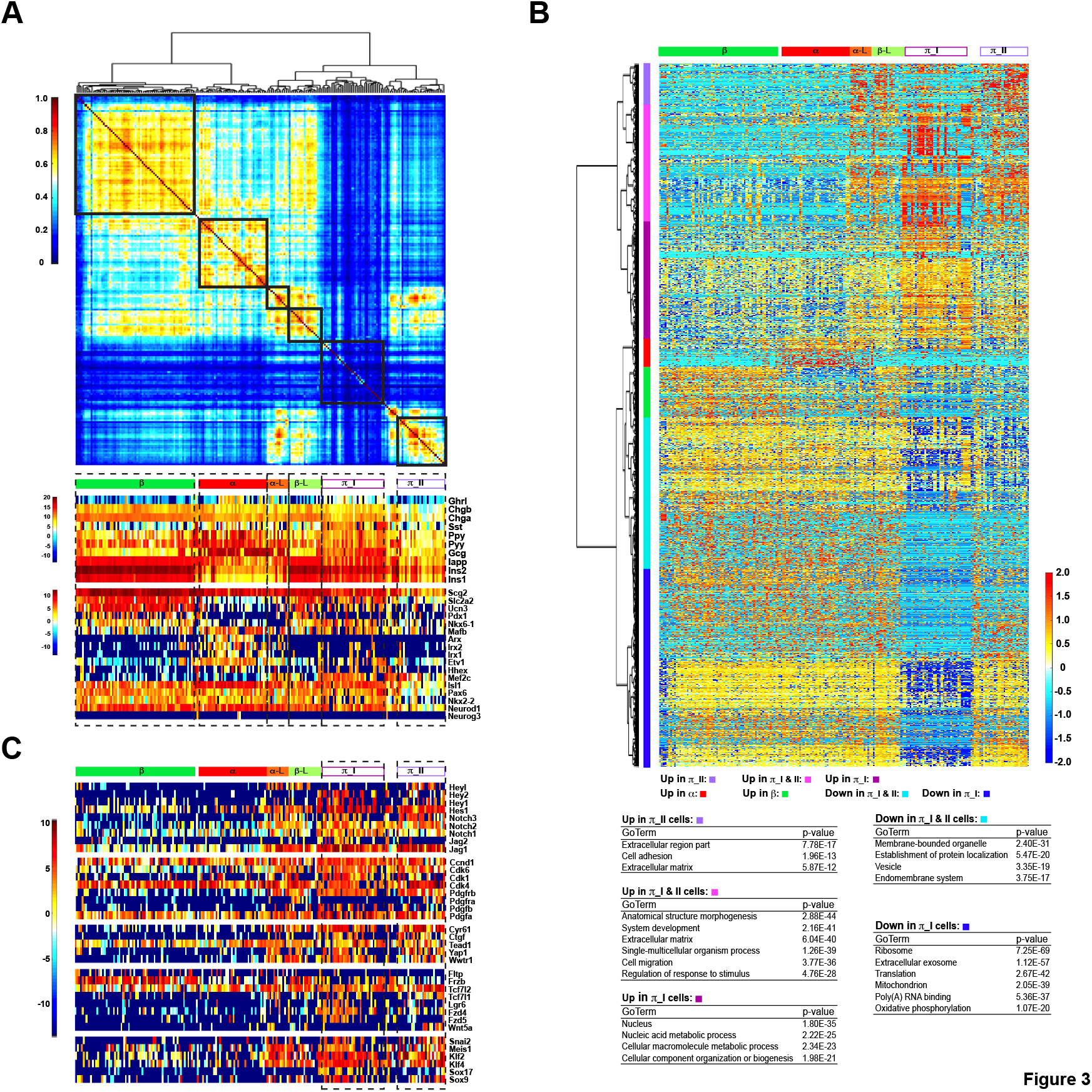
Single-cell transcriptome signature of P7.5 mouse islets (A) Up: Correlation coefficient cluster plot partitions P7.5 islet cells into β, α, β-like (β-L), α-like (α-L), and π_I and II subpopulations (three putative δ cells and four PP cells were identified based on the predominant expression of *Sst* or *Ppy*, respectively, compared to other hormones, but not enough to form separated clusters; seven cells between π_I and II groups were treated as unknown as Itgam was detected in these cells in addition to hormones). Color scale corresponds the absolute correlation coefficient of each pair of cells; Bottom: Heatmaps showing expression patterns of selected cell type specific hormones and transcription factors. Color scale corresponds the expression level (log_2_(RPKM+0.0001)). (B) Top: Heatmap showing 3911 differentially expressed genes (FDR < 0.01) of P7.5 islet single cells. Genes are clustered into 6 major groups, indicated by the colored bars next to the dendrogram and beneath the heatmap. Up in π_II: π_II enriched genes; Up in π_I and π_II: genes enriched in both π_I and π_II cells; Up in π_I: π_I enriched genes; Up in α: α cell enriched genes; Up in β: β cell enriched genes; Down in π_I and π_II: genes enriched in all cells except π_I and π_II cells; Down in π_I: genes enriched in all cells except π_I cells. Data (log_2_(RPKM+0.0001)) are standardized along each gene (row) such that the mean is 0 and the standard deviation is 1. Color scale corresponds the relative expression level. Bottom: Selected, top significantly enriched GO terms of each cluster. (C) Heatmaps showing expression patterns of selected π cell enriched genes, including components of Notch, PDGF, Hippo and Wnt signaling pathways, as well as several well-known progenitor associated genes. Color scale corresponds the expression level (log_2_(RPKM+0.0001)).

In addition to α and β cells, a substantial number of cells exhibited a gene expression pattern similar to π cells, and co-expressed transcription factors generally restricted to either α, β or δ cells in the adult islet (Fig. 3A). The P7.5 π cells can be divided into two sub-clusters, with π_I cells exhibiting somewhat higher, although still low, levels of hormone transcript (log_2_(RPKM+0.0001) = 5-10), while π_II cells exhibit even lower expression (log_2_(RPKM+0.0001) = 2-5) of hormone transcripts (Fig. 3A). As evident by the correlation matrix, π_I cells are molecularly diverse with pairwise correlation coefficients < 0.4, akin to adult π-cells (Fig. 2A, 3A). By comparison, π_II cells are relatively more homogenous. To exclude the possibility that π cells represent contaminating cells not being removed during islet purification, we examined the expression patterns of multiple alternative lineage-specific genes, and concluded that π cells were a distinct islet cell type (Supplemental Fig. S2B, S3A).

We also identified minor β cell and α cell subpopulations, indicating a greater degree of heterogeneity in cell identity during postnatal development. These cells display gene expression patterns that likely reflect transitional states and statistically cluster between the larger α cell and β cell populations and π population (Fig. 3A, B). We provisionally designated the minor subpopulation of β cells as β-like cells and, likewise, the minor population of α cells as α-like cells. Due to the relatively low number of cells in these two groups, they were excluded from downstream statistical analyses such as differential expression and pseudotime ordering (Materials and Methods).

### Development-associated genes and pathways are upregulated in P7.5 π cells

Differential expression analysis of cells isolated from P7.5 islets identified 3,911 genes. Hierarchical clustering partitioned these genes into seven modules: (1) α enriched genes, (2) β enriched genes, (3) π_I enriched genes, (4) π_II enriched genes, (5) genes enriched in both π_I and π_II cells, (6) genes enriched in all except π_I cells, and (7) genes enriched in all except π_I and π_II cells (Fig. 3B; Supplemental Table S3). Not surprisingly, the genes enriched in P7.5 β or α clusters are annotated to canonical β or α cell functions, such as vesicle formation and regulated hormone secretion. For instance, *G6pc2, Nkx6.1, Pcsk1, Pdx1, Slc2a*, and *Ucn3* were enriched in β cells, while Arx, *Irx1, Irx2*, and *Mafb* were enriched in α cells.

In contrast, π_I and π II cells were enriched for genes related to proliferative expansion of the endocrine islet, including genes related to diverse nuclear structure and biosynthesis (such as *Anapc1, E2f5, Mdm2, Plk2, Top1, Top2a*, and *Top2b*), extracellular matrix formation, cell adhesion, and cell migration, suggesting these cells might have roles in proliferation, endocrine islet tissue structure, and β cell maturation (Arous and Wehrle-Haller 2017). Furthermore, genes involved in stem cell/progenitor maintenance and tissue development were also enriched in π cells, including Krüppel-like factors *(Klf2, Klf4, Klf6, Klf12*, and *Klf14)* and Sry-related HMG box genes *(Sox4, Sox6, Sox9, Sox11* and *Sox17)*, as well as multiple signaling pathways that are known to govern developmental processes (Fig. 3B, C; Supplement Fig. S3A; Supplemental Table S3).

Notably, we identified significant upregulation of both Pdgf ligands and receptors in π_I and π_II cells. Given the established role for PDGF signaling in age-dependent β cell proliferation (Chen et al. 2011), this supports the interpretation that π cell populations have a critical role in the postnatal expansion of β cell mass. The expression pattern of PDGF pathway components implies that growth regulation involves both autocrine and paracrine mechanisms. For example, *Pdgfrb* and *Pdgfb* are upregulated in π_I and π_II cells, while *Pdgfra* is only upregulated in π_II cells, and *Pdgfa* was detected in majority of P7.5 islet cells (Fig. 3C). The upregulation of cell cycle genes such as *Cdk1, Cdk4, Cdk6* and *Ccnd1*, have been shown to induce β cell proliferation *in vitro* and *in vivo* and enhance human β cell engraftment (Vasavada et al. 2006; Fiaschi-Taesch et al. 2009; Fiaschi-Taesch et al. 2010; Tiwari et al. 2015). Consistent with the role of PDGF signaling in cell cycle activation (Chen et al. 2011), we found that *Cdk1* and *Cdk6*, as well as *Cdk11b, Cdk13* and *Cdk14* were enriched in the π_I and π_II cell populations, while *Ccnd1* and *Cdk4* were expressed in the majority of P7.5 islet cells (Fig. 3C).

In addition to PDGF signaling, π_I and π_II cells exhibited significant enrichment of Hippo and Wnt pathway components (Fig. 3C). These pathways have important roles in growth regulation and differentiation in many tissues, including pancreatic islet (Liu and Habener 2010; Shu et al. 2012; Bernal-Mizrachi et al. 2014; Takamoto et al. 2014; Cebola et al. 2015; George et al. 2015; Riley et al. 2015), however specific roles in postnatal β cell expansion have not been clearly defined. While Hippo-signaling effector gene *Tead1* was expressed in most P7.5 islet cells, the expression of its co-activators *Yapl* and *Wwtrl (Taz)* was limited to π_I and π_II cells. Importantly, two YAP1/WWTR1 downstream targets *Ctgf* and *Cyr61* were also enriched in π_I and π_II cells, suggesting YAP1 and WWTR1 are functionally active in π cells (Fig. 3C). We also observed differential expression of multiple components of the Wnt signaling cascade in P7.5 islet cells. While *Wnt4* was broadly expressed in the majority of P7.5 islet cells, *Wnt5a* was enriched in π_II cells, and *Fzd4, Fzd5, Tcf7l1* and *Tcf7l2* were enriched in both π_I and π_II cells. *Lgr6*, an R-spondin receptor gene and putative stem/progenitor cell marker, was also enriched in both π_I and π_II cells. In contrast, β cells exhibited upregulated expression of Wnt-antagonist *Frzb* and roughly half of the β cell population expressed *Fltp* (*l700009Pl7Rik*), a Wnt/PCP effector and newly discovered biomarker for mature β cells (Bader et al. 2016). Interestingly, *Fltp* was essentially silent in both π_I and π_II cells, suggesting that π cells are functionally immature (Fig. 3C; Supplemental Table S3).

Much to our surprise, Notch pathway ligands, receptors, and downstream targets were also dynamically upregulated in π_I and π_II cells. For instance, *Jag1, Notch3*, and the Notch downstream targets *Hes1, Hes3, Hey1*, and *Hey2* were broadly upregulated in both π populations. In contrast, *Jag2, Notch1*, and *Notch2* were primarily enriched in π_I cells, while *Heyl* was only upregulated in π_II cells (Fig. 3C, Supplemental Table S3).

### Comparative analysis of neonatal and young adult π cells

Multiple lines of evidence demonstrate that π cells cannot be explained as an artefactual result generated by cell doublets of different islet cell lineages. First, cell doublets tend to have more mRNA and generate more detectable genes than any single cell, while π cells generally have a lower or average number of detectable genes among all of the single cells (Supplemental Fig. S2C, S3B). Second, π cells display a unique transcriptional profile that cannot be explained by a combination of β, α or δ cells, or being dying cells, as demonstrated by the hierarchical clustering of differentially expressed genes (Fig. 2C; Supplemental Fig. S2A, S3B). For example, *Ghrl*, a hormone mainly associated with scarce ɛ cells in pancreatic islets, was enriched in both adult and P7.5 π cells, but remained undetectable in adult β, α, δ or PP cells (Fig. 2A). Even more compellingly, a substantial number of genes were distinctively silent or downregulated in π cells (in particular, P7.5 π_I and adult π cells) relative to α cells and β cells in either P7.5 or adult islets. This pattern directly supports the conclusion that π cells are a unique cell type. GO analysis revealed that the down-regulated genes are enriched in the processes of protein synthesis (including RPL and RPS gene family members, translation initiation and elongation factors, and protein transport genes) and mitochondrial oxidative phosphorylation (including Cytochrome C oxidase subunits, NADH dehydrogenases, and ATPases) (Fig. 2C, 3B; Supplemental Tables S2, S3). Recent studies of stem/progenitor cell biosynthesis and metabolism have suggested that, to maintain a long-term self-renewal capacity and to minimize oxidative damage, stem cells, including adult quiescent stem cells, keep low levels of global protein synthesis and rely on glycolysis instead of mitochondrial oxidation for energy (Tsatmali et al. 2005; Simsek et al. 2010; Signer et al. 2014; Llorens-Bobadilla et al. 2015; Wanet et al. 2015; Blanco et al. 2016).

The majority of π cells in both P7.5 and adult islets expressed the endocrine markers *Isl1, Neurod1*, and *Pax6* at levels comparable to those observed in β, α or δ cells;, neither population however expresses the mature β cell marker *Fltp.* Interestingly, π cells were also characterized by promiscuous co-expression of transcription factors and cell markers whose expression is otherwise restricted to distinct, canonical islet cell types, a common feature of stem and progenitor cells that is thought to be associated with lineage plasticity (Hu et al. 1997; Brunskill et al. 2014; Llorens-Bobadilla et al. 2015; Nimmo et al. 2015; Kim et al. 2016).

These characteristics of pancreatic π cells indicate that, although relatively rare, they represent a group of immature insulin-expressing cells with expression signatures similar to stem or progenitor cells in other tissues, as suggested by previous studies (Seaberg et al. 2004; Smukler et al. 2011). This is further supported by studies demonstrating that β cell postnatal maturation, marked with robust glucose-stimulated insulin secretion and decreased proliferation, is associated with enhanced mitochondrial oxidative ATP production (Bader et al. 2016; Yoshihara et al. 2016). In contrast, reduced insulin production promotes β cell proliferation in a cell autonomous manner (Szabat et al. 2016).

To further investigate the molecular dynamics of π cells related to other pancreatic islet cells in a developmental content, we generated additional scRNA-seq datasets from E18.5, P1.5, and P4.5 islets. These datasets were pooled and categorized as pre-P7.5 cells for downstream analysis. Importantly, cells with transcriptome profiles similar to P7.5 π_I and π_II cells were also detected in the pre-P7.5 group (although the proportion of these cells was relatively smaller than the one in P7.5 islets). Next, a quantitative measure of cellular progression known as pseudotemporal ordering was used to define the cell populations within pre-P7.5, P7.5, and early adult. We utilized an unsupervised algorithm in Monocle that improves the temporal resolution during a dynamic biological process, such as development (Trapnell et al. 2014; Llorens-Bobadilla et al. 2015). This enabled us to order islet cells of different ages based on differentially expressed genes of scRNA-seq datasets from five ages (FDR < 0.01; Materials and Methods; Supplemental Table S4). Interestingly, this analysis assigned the earliest points on the pseudotime axis to the majority of π_I, π_II, and adult π cells. The remaining α cells, β cells, δ cells, and PP cells were broadly dispersed across later points along the pseudotime axis, regardless of ages.

We next examined the pseudotemporal kinetics of select genes associated with various biological functions (Fig. 4; Supplemental Fig. S4). In alignment with peak β mass expansion during postnatal period, the expression of genes associated with proliferation, cell division, and tissue development and anatomical morphogenesis were significantly more enriched in postnatal π cells than adult π cells (Fig. 4A; Supplemental Fig. S4A), consistent with the idea that adult π cells represent quiescent progenitor cells. While on the other hand, as anticipated, the expression of genes involved in endocrine cell maturation and function was correlated with increasing pseudotime (Supplemental Fig. S4A), although β cell specific transcription factor *Nkx6.1* was already expressed in the majority of π cells (both neonatal and adult) (Fig. 4B). Furthermore, we found unique enrichment of *Rbpj*, a key mediator of canonical Notch signaling that acts as a transcriptional repressor in the absence of NICD, in adult π cells (Supplemental Fig. S4B).

**Figure 4.**
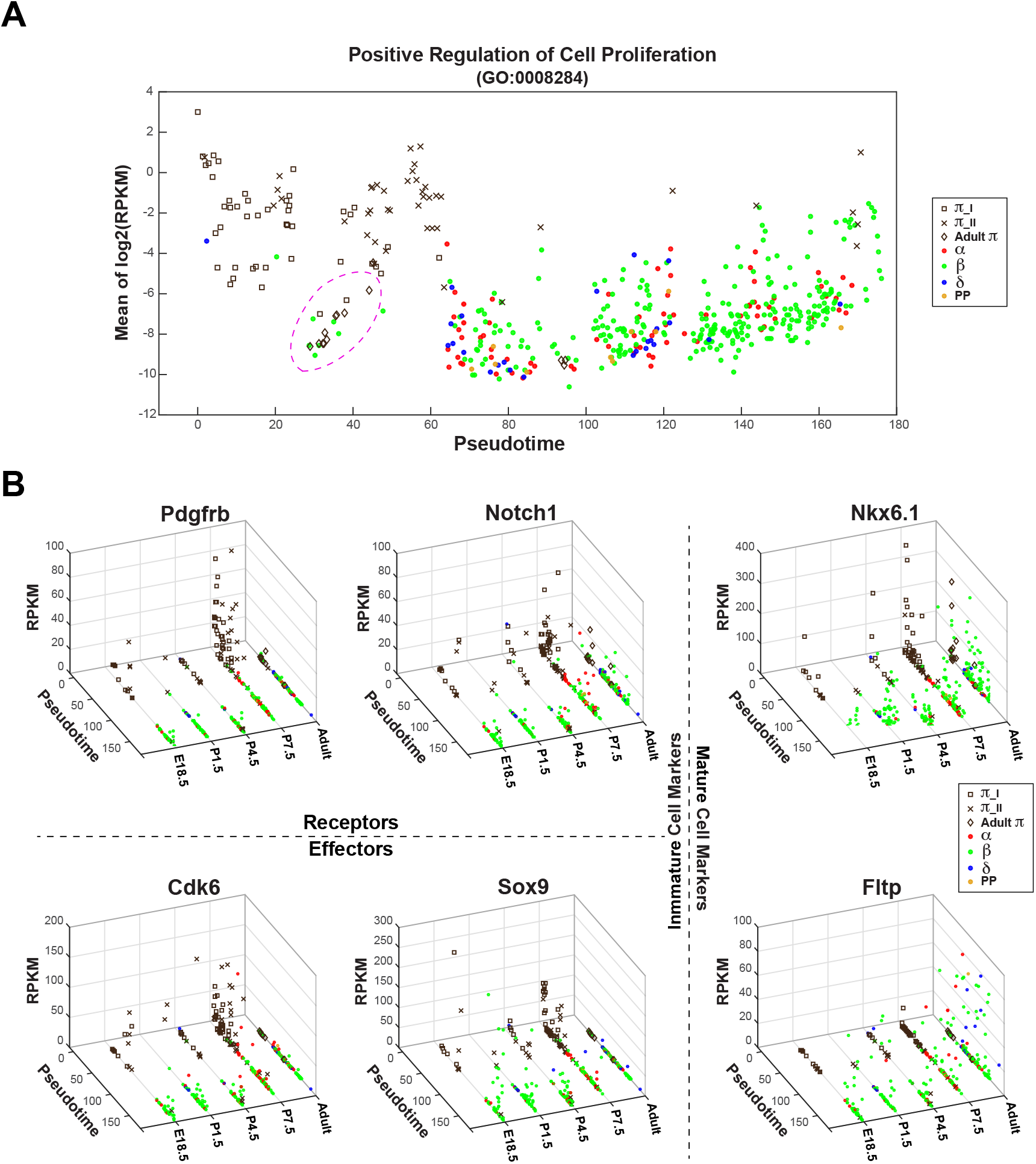
Developmental and pseutotemporal dynamics of gene expression in pancreatic islets (A) Expression pattern of selected, positive regulation of cell proliferation (GO:0008284) associated genes that were enriched in P7.5 π cells, as related to pseudotime. Islet cells pooled from five ages (E18.5, P1.5, P4.5, P7.5 and Adult) are ordered based on pseudotime (x-axis) and average expression levels (mean of log2(RPKM+0.0001)) of selected genes (y-axis). Each point represents a cell, and cell identity is indicated by color or shape. π_I and _II are π cells identified from E18.5, P1.5, P4.5 or P7.5. Majority of adult π cells are highlighted in dotted circle to show lacking of expression of cell proliferation associated genes. (B) Expression profiles of example genes *(Pdgfrb*, PDGF pathway target *Cdk6, Notch1*, Notch pathway target *Sox9*, β cell specific transcription factor *Nkx6.1*, and mature β cell marker *Fltp*), as related to pseudotime. Islet cells pooled from five ages are ordered based on pseudotime (x-axis), ages (y-axis) and expression level of the corresponding gene (RPKM).

### INS1^+^ PDGFRB^+^ P7.5 islet cells have higher proliferative potential

Single cell suspensions from purified P7.5 and young adult (10-week old) *MIP1-eGFP* islets were subjected to FACS analysis to determine the surface expression pattern of PDGFRB and NOTCH1 on GFP^+^ cells. As expected, much higher percentage of GFP^+^ P7.5 islet cells expressed PDGFRB or NOTCH1 or both on cell surface, compared to GFP^+^ young adult islet cells (Supplemental Fig. S5). To assess the proliferative potentials of P7.5 INS1^+^ subpopulations and to exclude the possibility that PDGFRB positive P7.5 islet cells are dying cells, we FACS purified the two GFP^+^ subpopulations, GFP^+^ PDGFRB^−^ and GFP^+^ PDGFRB+, and then measured EdU *in vitro* incorporation rate of each subpopulation with or without exposure to platelet-derived growth factor-A/B (PDGF-A/B), the ligands for PDGF signaling (Chen et al. 2017) (Materials and Methods). Remarkably, P7.5 GFP^+^ PDGFRB^+^ islet cells exhibited significantly higher proliferative potential than GFP^+^ PDGFRB^−^ islet cells, and this higher proliferation rate was amplified in the presence of PDGF ligands (two-way ANOVA, p < 0.0001), unlike GFP^+^ PDGFRB^−^ cells (unpaired t-test, p = 0.8206) (Fig. 5A; Materials and Methods). After 4-5 days of *in vitro* culture in the presence of PDGF-A/B, visible colonies were formed from GFP^+^ PDGFRB^+^ cells, as compared to GFP^+^ PDGFRB^−^ cells (Fig. 5A).

**Figure 5.**
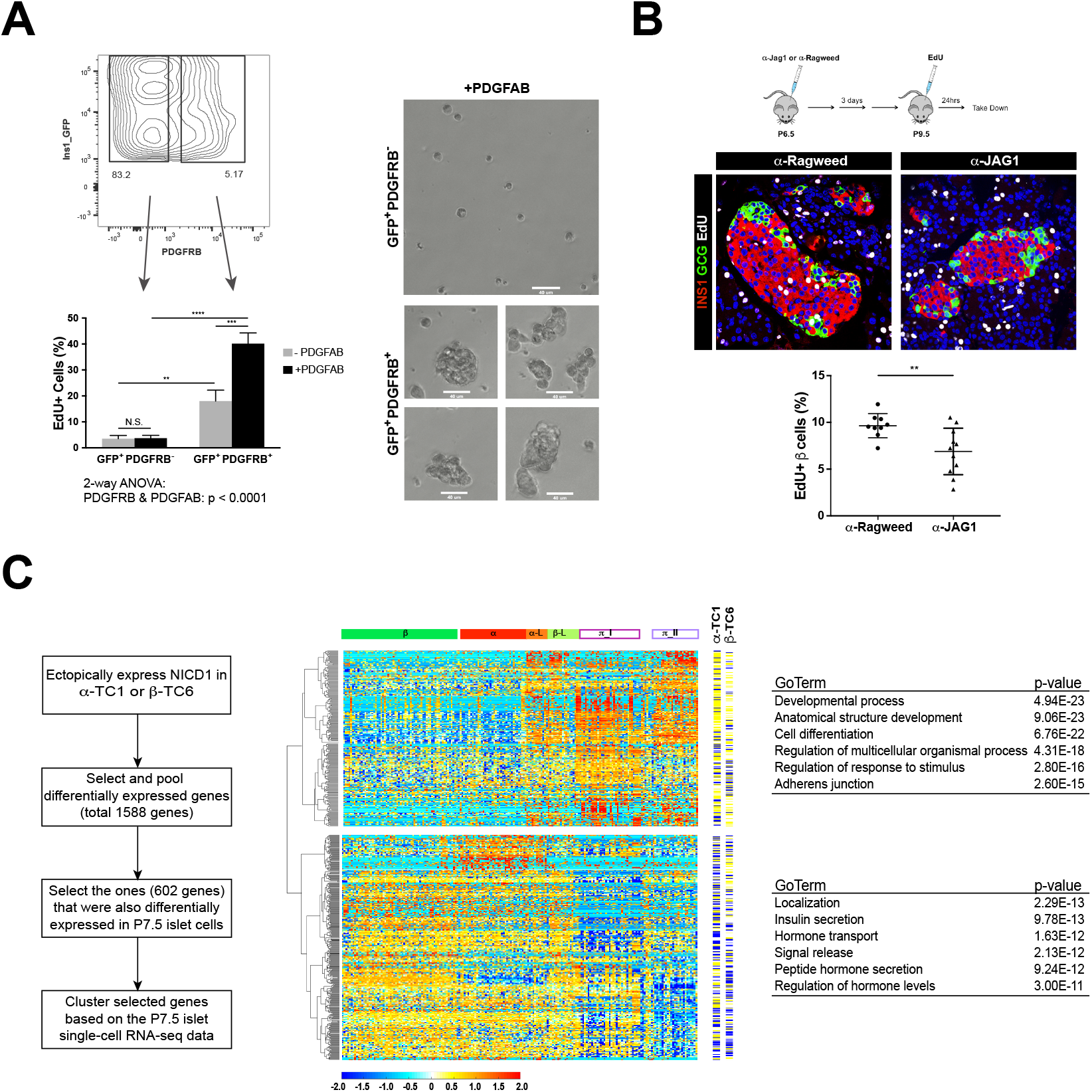
PDGF and Notch pathways are involved in regulating postnatal pancreatic endocrine proliferation and development. (A) Left: Ratios of EdU incorporation of FACS purified P7.5 *MIP1-e*GFP islet fractions (GFP^+^ PDGFRB^−^ or GFP^+^ PDGFRB+) with or without exposure to PDGF ligand (PDGFAB). Unpaired t-test: GFP^+^ PDGFRB^−^ vs. GFP^+^ PDGFRB+: p = 0.0046 (-PDGFAB) and p < 0.0001(+PDGFAB); with PDGFAB vs. without exposure to PDGFAB: p = 0.8206 (GFP^+^ PDGFRB^−^) and p = 0.0009 (GFP^+^ PDGFRB+). Two-way ANOVA test for interaction between PDGFRB and PDGFAB: p < 0.0001; Right: GFP^+^ PDGFRB^+^ cells formed visible colonies after 4~5 days of *in vitro* culture in the presence of PDGFAB, as compared to GFP^+^ PDGFRB^−^ under the same culture condition. Images for GFP^+^ PDGFRB^+^ *in vitro* culture (bottom) were taken from four different fields. Scale bar is at 40 um. (B) Notch blockade by JAG1 antagonistic antibody leads to significant reduction of pancreatic β cell proliferation in postnatal pups. Top: scheme of antibody treatment. Middle: representative immunofluorescence staining of islet from α-Ragweed (control) (left) and α-JAG1 (right) treated pups; Bottom: quantitative measurement of β cell proliferation rate (ratio of EdU incorporation of INS^+^ cells; 9 control pups vs. 11 α-JAG1 treated pups). (C) Left: scheme of *in vitro* assessing roles of NICD1 in endocrine lineage; Middle: Heatmap showing 602 NICD1 downstream targets in a-TC1-c9 or β-TC6 cell lines that are differentially expressed in P7.5 islet single cells. The yellow-blue color panels indicated whether individual genes are up (yellow) or downregulated (blue) in each corresponding cell line. Genes are clustered into two clusters: π cell enriched genes (top) and α and/or β cell enriched genes (bottom). Data (log_2_(RPKM+0.0001)) are standardized along each gene (row). Color scale corresponds the relative expression level. Right: Selected, top significantly enriched GO terms for each cluster.

### Notch is involved in postnatal β cell expansion and endocrine development

Notch signaling plays key roles in pancreatic development during embryogenesis (Shih et al. 2013; Bar and Efrat 2014); however, it is unclear how Notch activation affects endocrine development and β cell expansion after birth. Previous work has demonstrated a role for Notch pathway activation in the proliferation of cultured human β cells (Bar et al. 2008). Given the association of multiple Notch ligands, receptors and downstream targets with the proliferative populations in the postnatal islet, we sought to determine whether Notch signaling is required for postnatal β cell expansion.

While *Notch1, Notch2* and *Notch3* receptors exhibit complex overlapping expression in our scRNA-seq datasets, the Notch ligand *Jag1* is broadly expressed across the proliferative subsets of cells (Fig. 3C). Taking advantage of this, we treated postnatal mice (P6.5) with an antibody that antagonizes JAG1-dependent signaling (Lafkas et al. 2015), and assessed the proliferation of islet β cells by EdU labeling (Fig. 5B; Materials and Methods). JAG1 blockade significantly reduced the EdU incorporation rate in β cells (unpaired t-test: p = 0.0077), consistent with the hypothesis that JAG1-dependent regulation is required for maximal proliferative capacity of the postnatal progenitor population. Because Notch receptor expression is confined to the π_I and π_II cell populations at P7.5 this implies, as does the expression of PDGF receptors, that π cells are likely the source of postnatal β mass expansion.

To further explore possible roles for Notch signaling in the content of pancreatic endo-exocrine lineage determination, we ectopically expressed the intracellular domain of NOTCH1 (NICD1) in pancreas-derived β (βTC6), α cell (αTC1-c9) and acinar (AR42J) cell lines and evaluated changes in gene expression relative to the P7.5 scRNA-seq datasets we had already collected (Supplemental Table S5). In the acinar cell line, we observed reduced expression of the acinar markers *Ptf1a* and *Krt19* with a corresponding induction of the bipotent ductal/endocrine progenitor marker *Hnf1b.* In addition, multiple endocrine-related genes involved in hormone transport, along with several established Notch downstream targets, were significantly upregulated (Supplemental Fig. S5A). This suggests that active Notch signaling can direct cells toward an endocrine lineage fate.

The ectopic expression of NICD1 led to significant changes in the expression of 704 genes in βTC6 cells and 1,133 genes in αTC1-c9 cells. When compared to the differentially expressed genes in P7.5 islets, 602 of these NICD1-regulated genes were also differentially expressed in P7.5 islets (Supplemental Table S5). Hierarchical clustering of these genes using scRNA-seq data from the P7.5 islet cells revealed that a large proportion of genes upregulated by NICD1 in either αTC1-c9 cells, βTC6 cells or both cell lines exhibited increased expression in π cells relative to mature α cells and β cells. Gene ontology (GO) and signal pathway analysis indicates that these genes are generally associated with developmental tissue remodeling, consistent with the notion that these cells represent proliferative progenitors (Fig. 5C). Contrary to its previously defined role in ductal lineage identity, we did not observe NICD1-mediated upregulation of ductal specific genes such as *Bhlha15, Foxa2, Gata4, Hnflb, Krtl9, Mucl, Nr5a2, Onecut1, Pkd2*, and *Prox1* in either cell line (Supplemental Table S5).

Many of the genes downregulated by NICD1 in either αTC1-c9 cells or βTC6 cells are involved in the production and secretion of hormones, and showed a dynamic expression pattern in P7.5 islet cells, with decreased expression in π cells, indicating that the Notch pathway plays a role in keeping endocrine lineage cells in an immature state (Fig. 5C). This dampened hormone production and secretion is a hallmark of proliferative progenitor cells (Szabat et al. 2016) and further supports the conclusion that π cells in the young mouse drive postnatal expansion of the β cell compartment.

## Discussion

As the most significant period of new β cell formation in both humans and rodents, the postnatal period provides an ideal physiological setting for studying molecular mechanisms orchestrating functional β cell proliferation and differentiation (Meier et al. 2008; Chen et al. 2011; Blum et al. 2012; Arda et al. 2016; Bader et al. 2016). In the current study, we have profiled gene expression patterns of mouse islet cells at four neonatal time points (E18.5, P1.5, P4.5, and P7.5), as well as young adult (3-month old) by scRNA-seq. Based on global gene expression pattern of individual islet cells and profile of differentially expressed genes at each age, we uncovered a polyhormonal population (π cells) that is most evident and further separated into two subpopulations (π_I and π_II cells) in P7.5 islets when β cell mass is under rapid expansion. A substantial amount of genes involved in mitochondrial oxidative phosphorylation, global protein synthesis, and hormone transport and secretion are either silent or significantly downregulated in π cells of both neonatal and adult, indicating that π cells, although expressing much of transcription factors and hormones specific to mature β cells, are likely to be severely deficient in their ability to carry out glucose-stimulated insulin production and secretion. In contrast, postnatal π cells, but not adult π cells, are systematically enriched with genes involved in proliferation, tissue development, cellular component organization or biogenesis, and extracellular matrix formation, compared to other subpopulations in islets, consistent with their role in islet expansion during the postnatal period.

β cell mass postnatal expansion is regulated by PDGF signaling, however, it is unclear whether β cells share equal responsiveness to PDGF stimuli (Chen et al. 2011). Here, we found that PDGF pathway, accompanied by several downstream cell cycle genes, is mainly enriched in postnatal π cells. We further showed that in P7.5 islets INS1^+^ PDGFRB^+^ cells had a significantly higher proliferation potential, markedly enhanced by exposure to PDGF ligands, than PDGFRB^−^ INS1^+^ cells, which did not respond to PDGF stimuli. During embryonic development, the Notch pathway plays a pivotal role in promoting pancreatic progenitor proliferation and preventing premature endocrine differentiation (Bar and Efrat 2014). SOX9, a downstream target of Notch, is required for the adoption of pancreatic endocrine fate and maintains proliferation of the pancreatic endocrine progenitor pool (Seymour et al. 2007; Seymour et al. 2008). However, it remains elusive whether the Notch pathway is involved in postnatal β cell expansion. Like the PDGF pathway, the Notch pathway and downstream *Sox9* are also upregulated in postnatal π cells. We showed that acute blockade of JAG1 *in vivo* led to significant reduction in proliferative β cells during postnatal period, indicating the requirement for Notch pathway in β cell postnatal expansion. Ectopically expressed NICD1 in two islet derived cell lines activated genes involved in regulating developmental processes, while inhibited processes associated with hormone production and secretion. These data imply that while promoting cell proliferation, the Notch pathway also helps keep endocrine cells at an immature state during development. Interestingly, γ-secretase inhibitors, by blocking the Notch pathway, have been recently introduced into several protocols designed for *in vitro* generation of functional β cells from human pluripotent stem cell, applied at the last stages of the *in vitro* culture to promote β cell maturation (Pagliuca et al. 2014; Rezania et al. 2014; Massumi et al. 2016).

Reduction in mitochondrial oxidative phosphorylation and global protein synthesis are two defining features for stem cells and/or progenitors in several tissues or organs (Simsek et al. 2010; Llorens-Bobadilla et al. 2015; Wanet et al. 2015; Blanco et al. 2016). These are also characteristics of π cells in adult islets as well as π_I cells at P7.5. Although π_I and π_II cells share many molecular signatures and are both enriched with development-associated genes, there is intriguing difference between them. π_I cells are more depressed in global protein synthesis and mitochondrial phosphorylation, but more enriched with genes involved in nuclear structure and biogenesis, while π_II cells have greater expression of extracellular matrix and cell adhesion associated genes and their gene expression is more correlated with β-like cells. Pancreatic multipotent endocrine progenitors have been previously proposed to reside inside islets after birth (Seaberg et al. 2004; Smukler et al. 2011). Our findings suggest that postnatal π cells, activated by PDGF and Notch pathways, form a precursor pool supporting β cell postnatal proliferative expansion (Bar and Efrat 2014), and π_I, π_II and β-like cells represent sequential stages in the progression from a progenitor to an expanded β cell pool. By contrast, development associated signal pathways and genes are not upregulated in adult π cells, consistent with limited β cell regeneration in adult.

Two studies were recently published that used scRNA-seq to investigate the molecular mechanisms regulating postnatal β cell and/or α cell proliferation (Qiu et al. 2017; Zeng et al. 2017). In both studies, islet cells were pre-selected using FACS to enrich *mIns1-H2B-mCherry* positive β cells (Zeng et al. 2017), or *Ins1-RFP* positive β cells and *Gcg-Cre; Rosa-RFP* positive α cells (Qiu et al. 2017). Cells co-expressing multiple hormones were also excluded from data analysis (Qiu et al. 2017). In the current study, we apply an unbiased approach to survey islet cells at different ages, therefore include the π cells, which would predict to be excluded from stringent FACS purification due to the relatively lower hormone-specific promoter activity in these cells (judged by the generally lower expression levels of *Ins1* or *Gcg* in π cells compared to β or α cells, respectively; Supplemental Tables S2, S3). Our data demonstrates that postnatal β cell proliferation and maturation is a progression with multiple sequential stages, with π cell specific molecular features being gradually downregulated while β cell function associated genes being gradually upregulated. Indeed, in agreement with our data, Qiu et al. showed that postnatal β cells were more likely enriched with genes associated with developmental processes compared to adult β cells, while conversely genes involved in hormone production and secretion, were more likely upregulated in adult β cells than in postnatal β cells.

More studies are warranted to determine whether π cells are derived from differentiated β cells or, as suggested by their presence in the adult, exist as a separate progenitor stably persisting in islets. Also important will be to determine what the stimuli are that drive π cells from a quiescent state to a proliferative state in response to physiological demands on metabolism. Our data reveals the systematic molecular characteristics of the proliferative compartment driving postnatal expansion of β cells and provides the basis for addressing these and many other questions.

## Materials and Methods

### Animals, Islet Isolation, and Islet Dissociation

Animals used in this study included *MIP1-eGFP* (Hara 2003) and C57BL/6 mice (Charles River). All mice were housed under specific pathogen-free conditions. The Genentech Institutional Animal Care and Use Committee (IACUC) approved all animal studies. Due to the small and irregular shape of neonatal pancreatic islets, we isolated E18.5, P1.5, P4.5 and P7.5 islets from heterozygous *MIP1-eGFP* animals to minimize contamination from non-islet cell types. Young adult islets were collected from 10-12 week old C57BL/6 mice. E18.5, P1.5, P4.5, and P7.5 pancreatic tissue was dissected, cut into small pieces, and incubated in Liberase solution at 37°C for 10-15 minutes. P21 and adult pancreas tissues were perfused with Liberase solution through the common bile duct then dissected and incubated at 37°C for 10 minutes. Liberase solution was prepared by dissolving 5 mg Liberase TL (Roche, 05401020001) in 20 ml 1X HBSS buffer containing 25 mM HEPES (pH 7.2), 1.8 mM CaCl_2_, 10 μg/ml DNase, and 0.1% BSA. Islets were separated from acinar tissue using Histopaque 1077 (Sigma-Aldrich, 10771) density centrifugation. Purified islets were dissociated into single-cell suspension using Accumax cell dissociation solution (Innovative Cell Technologies, AM105) for 15-30 minutes at room temperature. For bulk RNA-seq studies, dissociated GFP^+^ β cells were isolated by fluorescence activated cell sorting (FACS) and dead cells were excluded using SYTOX Blue nucleic acid staining (ThermoFisher Scientific, S11348). For singlecell RNA-seq samples, islets were hand selected twice under a light microscope (young adult C57BL/6 islets) or ultraviolet microscope (E18.5, P1.5, P4.5 and P7.5 *MIPl-eGFP* islets) to minimize non-islet tissue contamination. Cells were dissociated with Accumax and single cells used for sequencing. Three or four biological replicates were collected at each age for bulk RNA-seq studies. One collection each of E18.5, P1.5, and P4.5 *MIPl-eGFP* islets, three independent collections of P7.5 *MIPl-eGFP* islets, and two independent collections of C57BL/6 young adult islets were used for single-cell RNA-seq studies.

### Cell Lines, Vectors, and Transfection

AR42J (CRL-1492), βTC6 (CRL-11506), and αTC1 clone 9 (αTC1-c9; CRL-2350) cell lines were obtained from ATCC and cultured according to ATCC protocols (https://www.atcc.org). cDNAs encoding the mouse *Notch1* intracellular domain (amino acids 1749-STOP) and IRES-eGFP were cloned in-frame into a pCAGGS-CMV vector to generate an NICD1-IRES-eGFP construct. Cells were transfected with NICD1-IRES-eGFP plasmid or IRES-eGFP control plasmid (1 μg of plasmid per 10^6^ cells) using an Amaxa Nucleofector (Lonza). Transfected cells were maintained in culture for 48 hours. GFP^+^ cells were FACS purified for RNA-seq studies. Since high NOTCH1 expression has been shown to promote the pancreatic ductal lineage, we selected transfected cells with an intermediate level of GFP intensity during FACS purification.

### Anti-JAG1 Treatment

P6.5 C57BL/6 pups were injected once intraperitoneally with anti-JAG1.b70 blocking antibodies at 15 mg/kg (Lafkas et al. 2015). Anti-Ragweed was used as nontargeting control at the same dose. Three days after antibody treatment, animals were injected intraperitoneally with EdU at 100 mg/kg (ThermoFisher Scientific, A10044). Animals were sacrificed 24 hours later, pancreata dissected, and tissues fixed in 10% neutralized formalin overnight at 4°C for histological analysis.

### Immunofluorescence, Microscopy, Flow Cytometry, and Islet Cell Culture

To evaluate β cell proliferation, mouse pancreata were dissected, fixed with 10% buffered formalin overnight at 4°C, and transferred to 70% ethanol. Dehydrated tissue was processed in a Tissue-Tek VIP overnight and embedded in paraffin the following day. The tissues were cut on a microtome at 5 μm and immunofluorescence staining was performed as previously described (Solloway et al. 2015). The following antibodies were used: guinea pig anti-Insulin (Abcam, ab7842), mouse anti-Glucagon (Sigma-Aldrich, G2654), and rabbit anti-Ki67 (Abcam, ab16667). EdU incorporated cells were stained using a Click-iT EdU kit (ThermoFisher Scientific, C10340), and nuclei were counterstained with Hoechst 33342 (ThermoFisher Scientific, H21492). Images were captured at 40X magnification using a Leica SPE confocal microscope. Approximately 3-4 sections (each separated by roughly 45 μm or ~9 sections) were quantified including 1,000-2,000 β cells (15-25 islet sections) were manually counted for each pancreas. Rat anti-PDGFRB (APA5) APC conjugated (ThermoFisher Scientific, 17-1401-81) and rabbit anti-NOTCH1 (D6F11) PE conjugated (Cell Signaling, 15004S) antibodies were used for staining *MIP1-eGFP* islet cells for FACS analysis or purification. Dead cells were excluded using SYTOX Blue nucleic acid staining (ThermoFisher Scientific, S11348). FACS purified GFP^+^ β cell fractions were cultured in a 96-well plate (7,500 cells per well), with or without PDGF-A/B (50 ng/ml; ThermoFisher Scientific, PHG0134) in DMEM/F12 (ThermoFisher Scientific, 11330032) medium containing 1X B27 Supplement (ThermoFisher Scientific, 12587010), 1X N2 Supplement (R&D Systems, AR003), 20 ng/ml EGF (Sigma-Aldrich, E9644), 10 ng/ml FGF2 (Sigma-Aldrich, F0291) and 2 ng/ml heparin (Sigma Aldrich, H4784). After 24 hours, cells were treated with 25μM EdU. Following an additional 24 hour incubation, cells were fixed with 3.7% formaldehyde and stained for EdU. Hoechst and EdU labeled cells were imaged using an ImageXpress Micro Widefield High-Content System (Molecular Devices, Sunnyvale, CA) fitted with a 20X Plan Fluor ELWD air immersion objective (0.45 NA) and a Photometrics Coolsnap HQ2 CCD camera. Automated multi-well imaging was performed using MetaXpress (version 5.1) software with laser detection and autofocus of cells located on the bottom surface of the well. For each well, 100 fields (10×10 grid, 30 μm spacing) were acquired and EdU^+^ nuclei were quantified using a MultiWavelength Cell Scoring image analysis module in MetaXpress software. An unpaired t-test and two-way ANOVA analysis of EdU incorporation rates were performed using Prism7. *In vitro* colonies were imaged using ancEVOS Cell Imaging System (ThermoFisher Scientific).

### Bulk RNA Library Preparation and Sequencing

RNA was extracted using an RNeasy micro kit (Qiagen) per the manufacturer’s protocol including an on-column DNase digestion. Total RNA was used for library generation using an Ovation RNA-Seq System v2 (NuGEN) and the libraries were barcoded using Ovation Rapid DR Multiplex System (NuGEN). Library size was confirmed using a 2100 Bioanalyzer (Agilent Technologies) and DNA concentration determined using a qPCR-based library quantification kit (KAPA). The libraries were multiplexed and sequenced on a HiSeq 2500 (Illumina) to generate 20-30 million singleend 50-base pair reads per library.

### Single-Cell RNA Library Preparation and Sequencing

Single-cell suspensions prepared from pancreatic islets were used to capture single cells on 10-17 μm (medium-size) integrated fluidic circuit chips using a C1 Single Cell Autoprep System per the manufacturer’s protocol (Fluidigm). Approximately 350,000 cells per ml were used to prepare the cell suspension at a 60:40 ratio of cells to the C1 suspension reagent prior to loading the chip. After the capture step, chips were visually examined on an EVOS microscope to exclude multiple cells and empty capture sites from library preparation. cDNAs were generated on-chip using a SMARTer Ultra™ Low RNA Kit (Clontech) for the Fluidigm C1 system. Sequencing libraries were prepared on 96-well plates using a Nextera XT DNA sample preparation kit (Illumina). Library size was confirmed using a 2100 Bioanalyzer (Agilent Technologies) and DNA concentration determined using a qPCR-based KAPA library quantification kit (Roche). All libraries were sequenced on a HiSeq 2500 platform to generate an average of 2-4 million single-end 50-bp reads per sample (Chen et al. 2017).

### RNA-seq Data Analysis

#### RNA-seq Data Processing

Ribosomal RNA reads were removed from RNA-seq datasets and the remaining reads were aligned to the mouse reference genome (NCBI Build 37) using GSNAP version ‘2013-10-10’. We allowed a maximum of two mismatches per 50 base sequence (parameters: ‘ -M 2 -n 10 -B 2 -i 1 -N 1 -w 200000 -E 1 –pairmax-rna=200000 –clip-overlap’). Transcript annotation was based on the Ensembl gene database (release 67). To quantify gene expression levels, the number of reads mapped to the exons of each gene was calculated (Liu et al. 2014).

#### Bulk RNA-seq

Differential expression between groups of bulk RNA-seq samples was computed as normalized counts using the R DESeq2 package. For FACS-isolated *MIP1-GFP* β cells, DESeq2 analysis yielded a total of 2,310 positive protein-coding genes (FDR < 0.01) between successive stages (E18.5 and P1.5, P1.5 and P7.5, P7.5 and P21, and P21 and adult) or between E18.5 and adult or P7.5 and adult. We defined these changes as differential expression during the β cell maturation process (Supplemental Table S1). To determine the overall temporal change in the molecular profile of these cells and the connection between different stages, we hierarchically clustered these 2,310 genes from all 16 samples along two dimensions with sample similarities clustered first (column) followed by genes (row) (based on log_2_(RPKM+0.0001))). Biological replicates were treated independently. Euclidean distance and Ward linkage were used (Figure 1C, MATLAB 8.4.0).

For NICD1 *in vitro* overexpression studies, DESeq2 analysis was performed using NICD1-expressing samples and empty-vector treated samples for each cell line. This analysis yielded 2,008 differentially expressed genes (NICD1 downstream target, FDR < 0.01) for AR42J, 1,133 for αTC1-c9, and 704 for βTC6 (Supplemental Table S5). *Single-Cell RNA-seq*

Since sequencing of unhealthy and dying cells tend to yield a lower number of detectable genes and doublets tend to yield a higher number of detectable genes, we filtered cells with unusually low or high numbers of reads (two times the standard deviation) out of our sequencing datasets. Therefore, of the total 593 single-cell RNA-seq datasets generated from the five islet stages (E18.5, P1.5, P4.5, P7.5, and adult), we retained 561 datasets for downstream analyses. Each dataset contained a set of 2,947 to 8,759 uniquely detected genes. The majority of noncoding RNAs were expressed at low levels below the threshold of detectability; therefore, we focused our analysis on the 24,391 protein-coding genes to minimize data noise (Supplemental Tables S2, S3 and S4).

To cluster cells based on expression similarity, we excluded genes with less than 0.1 RPKM in all 561 selected cells and genes that fell within in the bottom 10% variance bin. To unmask important cell subtypes or sub-states, we also excluded the top 15 highest expressed cell-type specific genes (genes that share similar expression patterns as *Ins1/2, Gcg* or *Sst*): *Akr1a1, Cox6a1, Fth1, Gcg, Hint1, Iapp, Ins1, Ins2, Ppy, Pyy, Sst, Tmed3, Ttr, Rn18s*, and *Snora81*, since these genes accounted for greater than 10% of the total mapped reads in the majority of datasets. Ultimately, 17,634 protein-coding genes passed our selection criteria. We next calculated the pairwise Pearson’s linear correlation coefficient between each pair of cells for individual age groups based on recalculated RPKM using the total number of counts for these 17,634 genes. E18.5, P1.5, P4.5 datasets were grouped as pre-P7.5, while P7.5 and adult remained independent groups. Hierarchical clustering of the resulting coefficient matrixes was performed using Euclidean distance and Ward linkage (MATLAB 8.4.0). Individual cells were then assigned an identity (α cell, β cell, δ cell, PP cell, α-like cell, β-like cell or π cell) based on the clustered correlation matrix and hormone expression profile (Figures 2A and 3A; Supplemental Tables S2.2, S3.2 and S4.). Five adult cells and seven P7.5 cells with unclear identity were labeled as unknown and excluded from further downstream analyses, since these cells formed isolated clusters and were enriched with Itgax or Itgam, or both.

To resolve molecular differences among different islet cell subpopulations at particular developmental stages, differential expression was computed using the R Monocle package (Trapnell et al. 2014). We evaluated genes with an expression level ≥ 0.1 RPKM in more than five analyzed cells. The analyses were only performed on cells that passed the aforementioned selections. Unknown cells were excluded from the adult population. Unknown cells, putative δ cells, PP cells, α-like, and β-like cells were also excluded from the P7.5 population to minimize noise. For P7.5 and adult datasets, we generated two differentially expressed gene lists (FDR < 0.01), one including all islet cells and the second including α and β cells (as well as δ and PP cells in adult) since many endocrine lineage-specific genes were also expressed at intermediate levels in π cells and tended to be underrepresented in the islet group analysis. The two lists were pooled together and defined as differentially expressed among single islet cells at either the P7.5 or adult stage. To examine the expression profile and identify cell type-specific markers, we hierarchically clustered single-cell RNA-seq data selected genes (log_2_(RPKM+0.0001)) for adult and P7.5 islet cells. Hierarchical clustering was performed on genes (row) using Euclidean distance and Ward linkage, while cells (column) were ordered based on the coefficient clustering matrix (Figures 2C, S2A, 3B, and 5C; MATLAB 8.4.0). Cell type-specific genes were selected based on hierarchical clustering and subjected to gene ontology and pathway analysis using gprofiler with default settings (http://biit.cs.ut.ee/gprofiler/index.cgi).

t-SNE (t-distributed stochastic neighbor embedding) analysis of adult islet single cells was performed on 154 cells (data points) with 1,327 differentially expressed genes (log_2_(RPKM+0.0001)) using the R package Rtsne (default setting), with an initial reduction of feature space by PCA (Figure 2B; https://github.com/jkrijthe/Rtsne; default settings).

To reconstruct the pseudotemporal ordering of various stage islet cells, single-cell RNA-seq data from cells of the three age groups were pooled together with α-like, β-like, and unknown cells excluded from the analysis. Monocle was used to perform differential expression analysis and significant genes (FDR < 0.01) were selected for “pseudo-time” ordering (num_path = 1, reverse = FALSE) (Trapnell et al. 2014).

To investigate whether the Notch pathway has a role in endocrine lineage identity, we combined 1,133 NICD1 target genes from αTC1-c9 datasets and 704 NICD1 target genes from βTC6 datasets to generate a master list of 1,588 unique NICD1 target genes. We compared this gene list to the list of genes that were differentially expressed in P7.5 islets. This comparison yielded 602 genes that were both NICD1 downstream targets in the endocrine cell lines and differentially expressed in P7.5 islets. To examine the expression pattern of NICD1 target genes in P7.5 islets, we hierarchically clustered P7.5 single-cell RNA-seq data of these 602 genes.

## Author contributions

A.S.P. and J.A.Z. conceived and designed the experiment. J.A.Z., C.G., D.S., M.K.B., N.K., H.N., and R.S. performed the experiments. J.A.Z. analyzed RNA-seq data. J.S and Z.M built cDNA libraries and performed NGS sequencing. All authors refined the manuscript.

---

**Figure S1:**
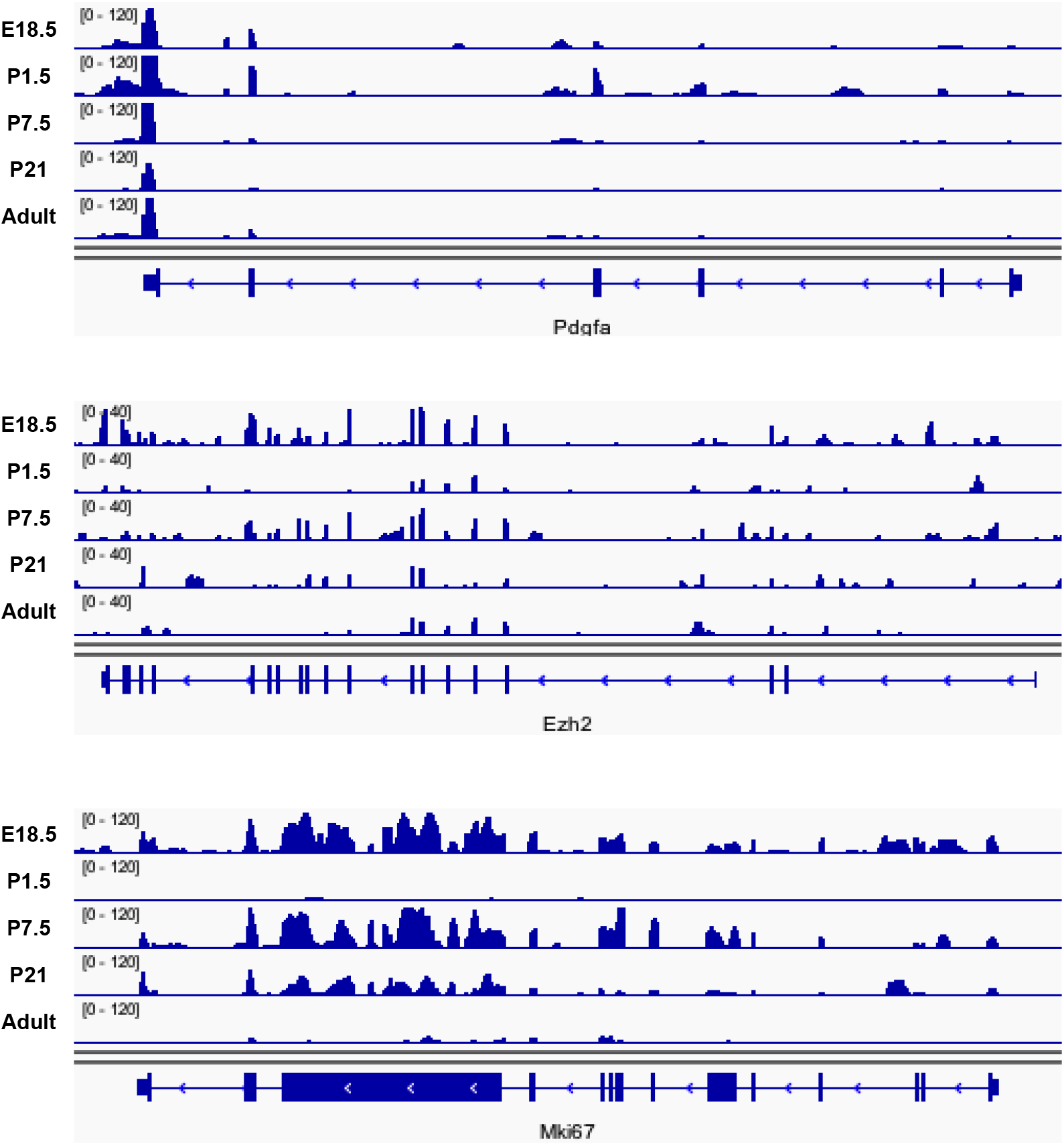
Expression patterns of *Pdgfa* (Top), *Ezh2* (middle) and *Mki67* (bottom) of FACS sorted *MIP1-GFP^+^* β cells across five ages: E18.5, P1.5, P7.5, P21 and adult. Datasets are presented in bigWig format using Integrative Genomics Viewer (IGV_2.3.83).

## References

Arda HE, Li L, Tsai J, Torre EA, Rosli Y, Peiris H, Spitale RC, Dai C, Gu X, Qu K et al. 2016. Age-Dependent Pancreatic Gene Regulation Reveals Mechanisms Governing Human beta Cell Function. Cell Metab 23: 909–920.

Arous C, Wehrle-Haller B. 2017. Role and impact of the extracellular matrix on integrin-mediated pancreatic beta-cell functions. Biol Cell.

Bader E, Migliorini A, Gegg M, Moruzzi N, Gerdes J, Roscioni SS, Bakhti M, Brandl E, Irmler M, Beckers J et al. 2016. Identification of proliferative and mature beta-cells in the islets of Langerhans. Nature 535: 430–434.

Bar Y, Efrat S. 2014. The NOTCH pathway in beta-cell growth and differentiation. Vitam Horm 95: 391–405.

Bar Y, Russ HA, Knoller S, Ouziel-Yahalom L, Efrat S. 2008. HES-1 is involved in adaptation of adult human beta-cells to proliferation in vitro. Diabetes 57: 2413–2420.

Bar Y, Russ HA, Sintov E, Anker-Kitai L, Knoller S, Efrat S. 2012. Redifferentiation of expanded human pancreatic beta-cell-derived cells by inhibition of the NOTCH pathway. J Biol Chem 287: 17269–17280.

Bernal-Mizrachi E, Kulkarni RN, Scott DK, Mauvais-Jarvis F, Stewart AF, Garcia-Ocana A. 2014. Human beta-cell proliferation and intracellular signaling part 2: still driving in the dark without a road map. Diabetes 63: 819–831.

Blanco S, Bandiera R, Popis M, Hussain S, Lombard P, Aleksic J, Sajini A, Tanna H, Cortes-Garrido R, Gkatza N et al. 2016. Stem cell function and stress response are controlled by protein synthesis. Nature 534: 335–340.

Blum B, Hrvatin SS, Schuetz C, Bonal C, Rezania A, Melton DA. 2012. Functional betacell maturation is marked by an increased glucose threshold and by expression of urocortin 3. Nat Biotechnol 30: 261–264.

Brennand K, Huangfu D, Melton D. 2007. All beta cells contribute equally to islet growth and maintenance. PLoS Biol 5: e163.

Brunskill EW, Park JS, Chung E, Chen F, Magella B, Potter SS. 2014. Single cell dissection of early kidney development: multilineage priming. Development 141: 3093–3101.

Butler PC, Meier JJ, Butler AE, Bhushan A. 2007. The replication of beta cells in normal physiology, in disease and for therapy. Nat Clin Pract Endocrinol Metab 3: 758–768.

Cebola I, Rodriguez-Segui SA, Cho CH, Bessa J, Rovira M, Luengo M, Chhatriwala M, Berry A, Ponsa-Cobas J, Maestro MA et al. 2015. TEAD and YAP regulate the enhancer network of human embryonic pancreatic progenitors. Nat Cell Biol 17: 615–626.

Chen H, Gu X, Liu Y, Wang J, Wirt SE, Bottino R, Schorle H, Sage J, Kim SK. 2011. PDGF signalling controls age-dependent proliferation in pancreatic beta-cells. Nature 478: 349–355.

Chen YJ, Friedman BA, Ha C, Durinck S, Liu J, Rubenstein JL, Seshagiri S, Modrusan Z. 2017. Single-cell RNA sequencing identifies distinct mouse medial ganglionic eminence cell types. Scientific reports 7: 45656.

Cheng CW, Villani V, Buono R, Wei M, Kumar S, Yilmaz OH, Cohen P, Sneddon JB, Perin L, Longo VD. 2017. Fasting-Mimicking Diet Promotes Ngn3-Driven betaCell Regeneration to Reverse Diabetes. Cell 168: 775–788 e712.

Chera S, Baronnier D, Ghila L, Cigliola V, Jensen JN, Gu G, Furuyama K, Thorel F, Gribble FM, Reimann F et al. 2014. Diabetes recovery by age-dependent conversion of pancreatic delta-cells into insulin producers. Nature 514: 503–507.

D’Amour KA, Bang AG, Eliazer S, Kelly OG, Agulnick AD, Smart NG, Moorman MA, Kroon E, Carpenter MK, Baetge EE. 2006. Production of pancreatic hormone-expressing endocrine cells from human embryonic stem cells. Nat Biotechnol 24: 1392–1401.

Dor Y, Brown J, Martinez OI, Melton DA. 2004. Adult pancreatic beta-cells are formed by self-duplication rather than stem-cell differentiation. Nature 429: 41–46.

Dorrell C, Schug J, Canaday PS, Russ HA, Tarlow BD, Grompe MT, Horton T, Hebrok M, Streeter PR, Kaestner KH et al. 2016. Human islets contain four distinct subtypes of beta cells. Nat Commun 7: 11756.

Fiaschi-Taesch N, Bigatel TA, Sicari B, Takane KK, Salim F, Velazquez-Garcia S, Harb G, Selk K, Cozar-Castellano I, Stewart AF. 2009. Survey of the human pancreatic beta-cell G1/S proteome reveals a potential therapeutic role for cdk-6 and cyclin D1 in enhancing human beta-cell replication and function in vivo. Diabetes 58: 882–893.

Fiaschi-Taesch NM, Salim F, Kleinberger J, Troxell R, Cozar-Castellano I, Selk K, Cherok E, Takane KK, Scott DK, Stewart AF. 2010. Induction of human beta-cell proliferation and engraftment using a single G1/S regulatory molecule, cdk6. Diabetes 59: 1926–1936.

George NM, Boerner BP, Mir SU, Guinn Z, Sarvetnick NE. 2015. Exploiting Expression of Hippo Effector, Yap, for Expansion of Functional Islet Mass. Mol Endocrinol 29: 1594–1607.

Gregg BE, Moore PC, Demozay D, Hall BA, Li M, Husain A, Wright AJ, Atkinson MA, Rhodes CJ. 2012. Formation of a human beta-cell population within pancreatic islets is set early in life. J Clin Endocrinol Metab 97: 3197–3206.

Gutierrez GD, Gromada J, Sussel L. 2017. Heterogeneity of the Pancreatic Beta Cell. Front Genet 8: 22.

Hara M, Wang X, Kawamura T, Bindokas VP, Dizon RF, Alcoser SY, Magnuson MA, Bell GI. 2003. Transgenic mice with green fluorescent protein-labeled pancreatic beta-cells. American journal of physiology Endocrinology and metabolism 284: E177–183.

Heit JJ, Karnik SK, Kim SK. 2006. Intrinsic regulators of pancreatic beta-cell proliferation. Annual review of cell and developmental biology 22: 311–338.

Hu M, Krause D, Greaves M, Sharkis S, Dexter M, Heyworth C, Enver T. 1997. Multilineage gene expression precedes commitment in the hemopoietic system. Genes & development 11: 774–785.

Kim TH, Saadatpour A, Guo G, Saxena M, Cavazza A, Desai N, Jadhav U, Jiang L, Rivera MN, Orkin SH et al. 2016. Single-Cell Transcript Profiles Reveal Multilineage Priming in Early Progenitors Derived from Lgr5(+) Intestinal Stem Cells. Cell Rep 16: 2053–2060.

Krishnamurthy J, Ramsey MR, Ligon KL, Torrice C, Koh A, Bonner-Weir S, Sharpless NE. 2006. p16INK4a induces an age-dependent decline in islet regenerative potential. Nature 443: 453–457.

Lafkas D, Shelton A, Chiu C, de Leon Boenig G, Chen Y, Stawicki SS, Siltanen C, Reichelt M, Zhou M, Wu X et al. 2015. Therapeutic antibodies reveal Notch control of transdifferentiation in the adult lung. Nature 528: 127–131.

Lenz A, Toren-Haritan G, Efrat S. 2014. Redifferentiation of adult human beta cells expanded in vitro by inhibition of the WNT pathway. PLoS One 9: e112914.

Linnemann AK, Baan M, Davis DB. 2014. Pancreatic beta-cell proliferation in obesity. Adv Nutr 5: 278–288.

Liu J, McCleland M, Stawiski EW, Gnad F, Mayba O, Haverty PM, Durinck S, Chen YJ, Klijn C, Jhunjhunwala S et al. 2014. Integrated exome and transcriptome sequencing reveals ZAK isoform usage in gastric cancer. Nature communications 5: 3830.

Liu Z, Habener JF. 2010. Wnt signaling in pancreatic islets. Advances in experimental medicine and biology 654: 391–419.

Llorens-Bobadilla E, Zhao S, Baser A, Saiz-Castro G, Zwadlo K, Martin-Villalba A. 2015. Single-Cell Transcriptomics Reveals a Population of Dormant Neural Stem Cells that Become Activated upon Brain Injury. Cell Stem Cell 17: 329–340.

Massumi M, Pourasgari F, Nalla A, Batchuluun B, Nagy K, Neely E, Gull R, Nagy A, Wheeler MB. 2016. An Abbreviated Protocol for In Vitro Generation of Functional Human Embryonic Stem Cell-Derived Beta-Like Cells. PLoS One 11: e0164457.

Meier JJ, Butler AE, Saisho Y, Monchamp T, Galasso R, Bhushan A, Rizza RA, Butler PC. 2008. Beta-cell replication is the primary mechanism subserving the postnatal expansion of beta-cell mass in humans. Diabetes 57: 1584–1594.

Nimmo RA, May GE, Enver T. 2015. Primed and ready: understanding lineage commitment through single cell analysis. Trends Cell Biol 25: 459–467.

Nir T, Melton DA, Dor Y. 2007. Recovery from diabetes in mice by beta cell regeneration. J Clin Invest 117: 2553–2561.

Pagliuca FW, Millman JR, Gurtler M, Segel M, Van Dervort A, Ryu JH, Peterson QP, Greiner D, Melton DA. 2014. Generation of functional human pancreatic beta cells in vitro. Cell 159: 428–439.

Petri A, Ahnfelt-Ronne J, Frederiksen KS, Edwards DG, Madsen D, Serup P, Fleckner J, Heller RS. 2006. The effect of neurogenin3 deficiency on pancreatic gene expression in embryonic mice. J Mol Endocrinol 37: 301–316.

Qiu WL, Zhang YW, Feng Y, Li LC, Yang L, Xu CR. 2017. Deciphering Pancreatic Islet beta Cell and alpha Cell Maturation Pathways and Characteristic Features at the Single-Cell Level. Cell Metab 25: 1194–1205 e1194.

Rahier J, Guiot Y, Goebbels RM, Sempoux C, Henquin JC. 2008. Pancreatic beta-cell mass in European subjects with type 2 diabetes. Diabetes Obes Metab 10 Suppl 4: 32–42.

Rezania A, Bruin JE, Arora P, Rubin A, Batushansky I, Asadi A, O’Dwyer S, Quiskamp N, Mojibian M, Albrecht T et al. 2014. Reversal of diabetes with insulin-producing cells derived in vitro from human pluripotent stem cells. Nat Biotechnol 32: 1121–1133.

Riley KG, Pasek RC, Maulis MF, Peek J, Thorel F, Brigstock DR, Herrera PL, Gannon M. 2015. Connective tissue growth factor modulates adult beta-cell maturity and proliferation to promote beta-cell regeneration in mice. Diabetes 64: 1284–1298.

Romer AI, Sussel L. 2015. Pancreatic islet cell development and regeneration. Current opinion in endocrinology, diabetes, and obesity 22: 255–264.

Russ HA, Ravassard P, Kerr-Conte J, Pattou F, Efrat S. 2009. Epithelial-mesenchymal transition in cells expanded in vitro from lineage-traced adult human pancreatic beta cells. PLoS One 4: e6417.

Seaberg RM, Smukler SR, Kieffer TJ, Enikolopov G, Asghar Z, Wheeler MB, Korbutt G, van der Kooy D. 2004. Clonal identification of multipotent precursors from adult mouse pancreas that generate neural and pancreatic lineages. Nat Biotechnol 22: 1115–1124.

Seymour PA, Freude KK, Dubois CL, Shih HP, Patel NA, Sander M. 2008. A dosage-dependent requirement for Sox9 in pancreatic endocrine cell formation. Dev Biol 323: 19–30.

Seymour PA, Freude KK, Tran MN, Mayes EE, Jensen J, Kist R, Scherer G, Sander M. 2007. SOX9 is required for maintenance of the pancreatic progenitor cell pool. Proc Natl Acad Sci U S A 104: 1865–1870.

Shih HP, Wang A, Sander M. 2013. Pancreas organogenesis: from lineage determination to morphogenesis. Annual review of cell and developmental biology 29: 81–105.

Shu L, Zien K, Gutjahr G, Oberholzer J, Pattou F, Kerr-Conte J, Maedler K. 2012. TCF7L2 promotes beta cell regeneration in human and mouse pancreas. Diabetologia 55: 3296–3307.

Signer RA, Magee JA, Salic A, Morrison SJ. 2014. Haematopoietic stem cells require a highly regulated protein synthesis rate. Nature 509: 49–54.

Simsek T, Kocabas F, Zheng J, Deberardinis RJ, Mahmoud AI, Olson EN, Schneider JW, Zhang CC, Sadek HA. 2010. The distinct metabolic profile of hematopoietic stem cells reflects their location in a hypoxic niche. Cell Stem Cell 7: 380–390.

Smukler SR, Arntfield ME, Razavi R, Bikopoulos G, Karpowicz P, Seaberg R, Dai F, Lee S, Ahrens R, Fraser PE et al. 2011. The adult mouse and human pancreas contain rare multipotent stem cells that express insulin. Cell Stem Cell 8: 281–293.

Solloway MJ, Madjidi A, Gu C, Eastham-Anderson J, Clarke HJ, Kljavin N, Zavala-Solorio J, Kates L, Friedman B, Brauer M et al. 2015. Glucagon Couples Hepatic Amino Acid Catabolism to mTOR-Dependent Regulation of alpha-Cell Mass. Cell reports 12: 495–510.

Szabat M, Page MM, Panzhinskiy E, Skovso S, Mojibian M, Fernandez-Tajes J, Bruin JE, Bround MJ, Lee JT, Xu EE et al. 2016. Reduced Insulin Production Relieves Endoplasmic Reticulum Stress and Induces beta Cell Proliferation. Cell Metab 23: 179–193.

Takamoto I, Kubota N, Nakaya K, Kumagai K, Hashimoto S, Kubota T, Inoue M, Kajiwara E, Katsuyama H, Obata A et al. 2014. TCF7L2 in mouse pancreatic beta cells plays a crucial role in glucose homeostasis by regulating beta cell mass. Diabetologia 57: 542–553.

Talchai C, Lin HV, Kitamura T, Accili D. 2009. Genetic and biochemical pathways of beta-cell failure in type 2 diabetes. Diabetes Obes Metab 11 Suppl 4: 38–45.

Teta M, Long SY, Wartschow LM, Rankin MM, Kushner JA. 2005. Very slow turnover of beta-cells in aged adult mice. Diabetes 54: 2557–2567.

Teta M, Rankin MM, Long SY, Stein GM, Kushner JA. 2007. Growth and regeneration of adult beta cells does not involve specialized progenitors. Dev Cell 12: 817–826.

Tiwari S, Roel C, Wills R, Casinelli G, Tanwir M, Takane KK, Fiaschi-Taesch NM. 2015. Early and Late G1/S Cyclins and Cdks Act Complementarily to Enhance Authentic Human beta-Cell Proliferation and Expansion. Diabetes 64: 3485–3498.

Trapnell C, Cacchiarelli D, Grimsby J, Pokharel P, Li S, Morse M, Lennon NJ, Livak KJ, Mikkelsen TS, Rinn JL. 2014. The dynamics and regulators of cell fate decisions are revealed by pseudotemporal ordering of single cells. Nature biotechnology 32: 381–386.

Tsatmali M, Walcott EC, Crossin KL. 2005. Newborn neurons acquire high levels of reactive oxygen species and increased mitochondrial proteins upon differentiation from progenitors. Brain Res 1040: 137–150.

Uchida T, Nakamura T, Hashimoto N, Matsuda T, Kotani K, Sakaue H, Kido Y, Hayashi Y, Nakayama KI, White MF et al. 2005. Deletion of Cdkn1b ameliorates hyperglycemia by maintaining compensatory hyperinsulinemia in diabetic mice. Nat Med 11: 175–182.

Vasavada RC, Gonzalez-Pertusa JA, Fujinaka Y, Fiaschi-Taesch N, Cozar-Castellano I, Garcia-Ocana A. 2006. Growth factors and beta cell replication. Int J Biochem Cell Biol 38: 931–950.

Wanet A, Arnould T, Najimi M, Renard P. 2015. Connecting Mitochondria, Metabolism, and Stem Cell Fate. Stem Cells Dev 24: 1957–1971.

Wang YJ, Golson ML, Schug J, Traum D, Liu C, Vivek K, Dorrell C, Naji A, Powers AC, Chang KM et al. 2016. Single-Cell Mass Cytometry Analysis of the Human Endocrine Pancreas. Cell Metab 24: 616–626.

Xin Y, Kim J, Ni M, Wei Y, Okamoto H, Lee J, Adler C, Cavino K, Murphy AJ, Yancopoulos GD et al. 2016. Use of the Fluidigm C1 platform for RNA sequencing of single mouse pancreatic islet cells. Proc Natl Acad Sci U S A 113: 3293–3298.

Yoshihara E, Wei Z, Lin CS, Fang S, Ahmadian M, Kida Y, Tseng T, Dai Y, Yu RT, Liddle C et al. 2016. ERRgamma Is Required for the Metabolic Maturation of Therapeutically Functional Glucose-Responsive beta Cells. Cell Metab 23: 622634.

Zeng C, Mulas F, Sui Y, Guan T, Miller N, Tan Y, Liu F, Jin W, Carrano AC, Huising MO et al. 2017. Pseudotemporal Ordering of Single Cells Reveals Metabolic Control of Postnatal beta Cell Proliferation. Cell Metab 25: 1160–1175 e1111.

Zhang J, McKenna LB, Bogue CW, Kaestner KH. 2014. The diabetes gene Hhex maintains delta-cell differentiation and islet function. Genes & development 28: 829–834.

